# Fur microbiome as a putative source of symbiotic bacteria in sucking lice

**DOI:** 10.1101/2024.04.01.587557

**Authors:** Jana Martin Říhová, Shruti Gupta, Eva Nováková, Václav Hypša

## Abstract

Symbiosis between insects and bacteria has been established countless times. While it is well known that the symbionts originated from a variety of different bacterial taxa, it is usually difficult to determine their environmental source and a route of their acquisition by the host. In this study, we address this question using a model of Neisseriaceae symbionts in rodent lice. These bacteria established their symbiosis independently with different louse taxa (*Polyplax, Hoplopleura, Neohaematopinus*), most likely from the same environmental source. We first applied amplicon analysis to screen for candidate source bacterium in the louse environment, that is, three species of rodents (*Microtus arvalis, Clethrionomys glareolus*, and *Apodemus flavicollis)*. The screened samples included rodent fur, skin, spleen, and ectoparasites sampled from the rodents. The amplicon analysis revealed a Neisseriaceae bacterium, closely related to the known louse symbionts. We assembled genome drafts of this environmental bacterium from all three rodent hosts. The sizes of the three drafts converged to a remarkably small size of approximately 1.4 Mbp, which is even smaller than the genomes of the related symbionts. Based on these findings, we propose a hypothetical scenario of the genome evolution during the transition of a free-living bacterium to the member of the rodent fur-associated microbiome and subsequently to the facultative and obligate louse symbionts.

## Introduction

Insects frequently establish symbiotic relationships with bacteria. Based on the evolutionary time elapsed and the host’s requirements (e.g., need for compounds missing in the diet), these symbioses undergo evolution throughout different evolutionary stages, potentially leading to obligate mutualism (McCutcheon et al 2019). The evolution from nascent symbionts (often called facultative symbionts) to obligate mutualists has been extensively investigated and many typical features of this process have been described. Less known, however, are the circumstances underlying the initial steps of symbiosis establishment. To propose a meaningful hypothesis on the origin of particular symbiotic relationships, one needs to know which free-living bacteria are the closest relatives of the symbiont, and what features of these bacteria may have played a role in the symbiosis establishment. Unfortunately, the precise phylogenetic origin of many symbionts remains uncertain. This is mainly due to dramatic changes in the genomes of obligate symbionts. In phylogenetic reconstructions such bacteria often form long branches with uncertain and unstable position in phylogenetic trees (Husnik et al 2011). Another frequent problem is a big evolutionary “gap” between the symbiont and its known free-living relatives. Specifically, there is a lack of intermediate stages in the evolutionary process, whose lifestyle would suggest a particular route to endosymbiosis. A rare example of phylogenetic pattern and lifestyle as a background for a symbiosis-origin hypothesis is the close relationship between the tick symbionts and pathogenic tick-borne bacteria: it has been hypothesized that at least in some cases the symbionts may have evolved from a pathogen transmitted by ticks to vertebrates (Gerhart et al 2016). A similar example of close relationships between pathogens and symbionts provides the opportunistic human infective *Sodalis praecaptivus*, related to the rich cluster of insect symbionts (Chari et al 2015). However, while allowing for genomic/functional comparison of bacteria in different stages of evolution towards symbiosis, this system does not provide any direct information on the origin (i.e. the source) of the symbionts.

In this study, we investigate a possible relationship between the symbiotic bacteria in rodent lice (*Polyplax, Hoplopleura*) and fur microbiomes of the rodent hosts. This model was chosen for several reasons. First, lice (Anoplura) are permanent ectoparasites of mammals, living in all developmental stages in the host fur. Their environment is thus stable, well defined, and easy to examine for its microbiome composition. Second, since lice feed exclusively on the vertebrate blood, they belong to the insect groups which require symbiotic bacteria as source of the essential compounds missing in their diet (typically B vitamins; Husnik 2018). However, compared to other insects, anoplurans are notable for high taxonomic diversity of the symbionts, due to frequent processes of the symbiont’s acquisitions and replacements. As a result, different groups of Anoplura host phylogenetically distinct, and often very distant bacterial taxa. Third, recent studies revealed that within this diversity, one bacterial lineage related to Neisseriaceae established symbiosis independently with several groups of lice and reached different stages of evolution. While in some lice they show features typical for facultative symbionts (e.g., *Polyplax serrata, Neohaematopinus pacificus*, and possibly *Haematopinus apri*; Nishide et al 2022, Rihova et al 2022, Rihova et al 2021), in others they function as obligate mutualists (*Hoplopleura acanthopus*; Rihova et al 2021). This frequent occurrence of the Neisseriaceae-related symbionts suggests that they may be acquired from the host, either as part of the fur/skin microbiome or from the host blood. Using amplicon screening, genomic comparisons, and phylogenetic reconstruction, we show that bacteria closely related to the symbionts are indeed frequent constituents of the host fur microbiome.

### Results and discussion

### Amplicon-based screening

The amplicon sequencing of the 16S rRNA gene libraries generated high quality amplicon data. The profiles of positive controls (staggered and even mock communities) were complete and confirmed high enough resolution, recovering rare mock taxa (0.04% relative abundance). The screening of environmental microbiomes associated with three rodent species (*Microtus arvalis, Clethrionomys glareolus*, and *Apodemus flavicollis*) indicates that their fur and skin contain bacteria which may potentially become a source of the typical symbionts in lice and other ectoparasites. More specifically, the microbiomes obtained from three different sources (fur swabs, skin and ectoparasites) contained several OTUs assigned to Neisseriaceae and related to the symbionts previously reported from rodent lice (Fig 1; complete overview of the microbiome profiles is provided in the Supplementary Table 1). Moreover, two of these OTUs (OTU 5 and OTU 815) were particularly abundant, in some samples exceeding 50% of total reads (and OTU 5 even reaching 97% in the *M. arvalis* fur swab; Fig. 1 and Supplementary Table 1). The absence of these OTUs in the rodent spleen (Supplementary Table 1) indicates that they are not vertebrate pathogens but in the rodent host they only inhabit fur and skin. On the other hand, their occasional occurrence in the ectoparasite sample is most likely due to surface contamination obtained from the host fur. The third abundant bacterium, OTU 66 was only isolated from the louse *Hoplopleura acanthopus* (sample 2MA from *M. arvalis*) and it corresponds to the previously described Neisseriaceae-related symbiont of *H. acanthopus* (Rihova et al 2021; Fig. 1). In the following text, when discussing individual specimens of the dominant OTU 5 from *Microtus arvalis, Apodemus flavicollis*, and *Clethrionomys glareolus*, we use names OTU5-Ma, OTU5-Af, and OTU5-Cg, respectively.

**Table 1.**
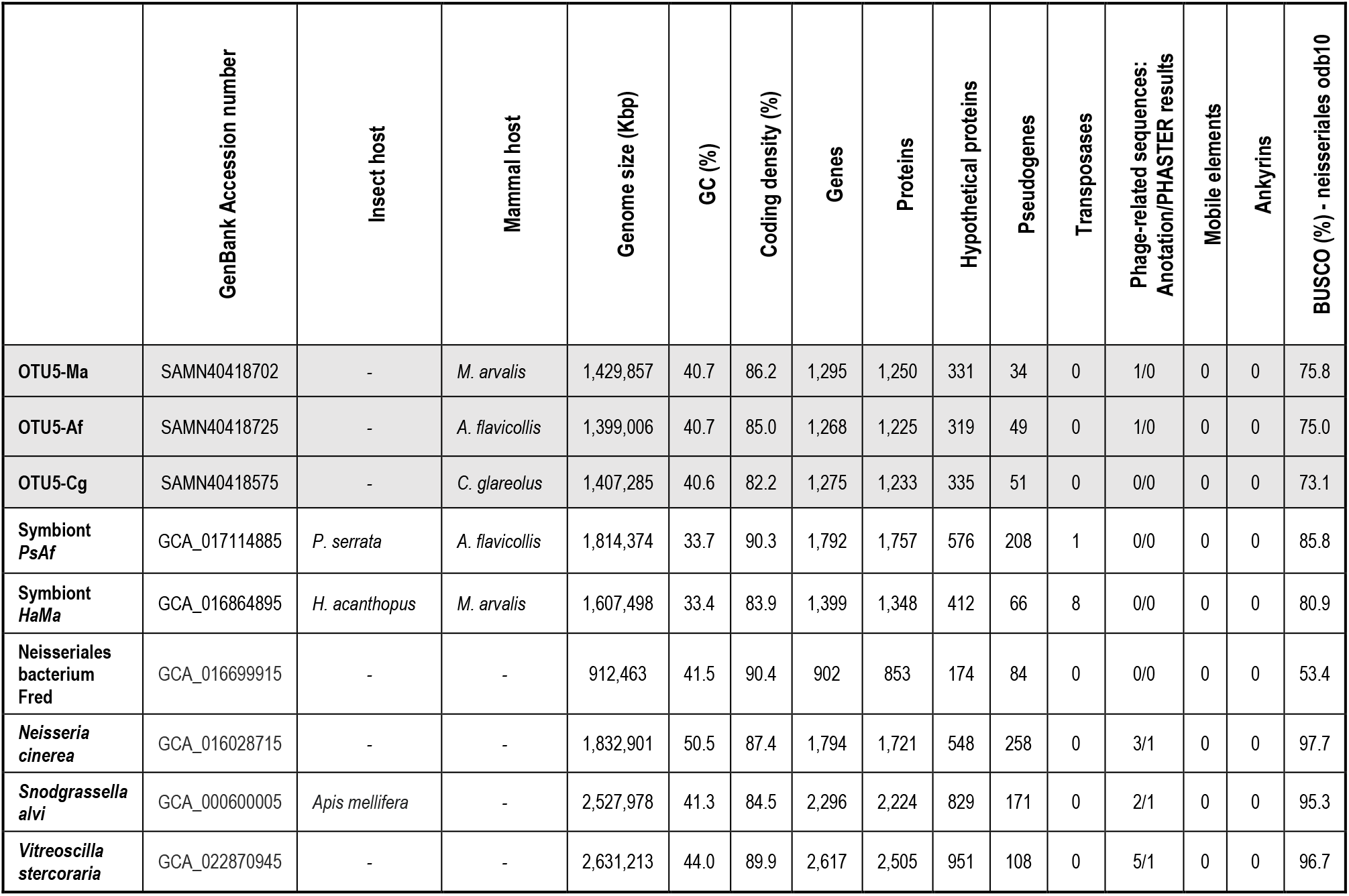
Comparison of the main genome characteristics of the new OTU 5 bacteria and their relatives.

**Figure 1.**
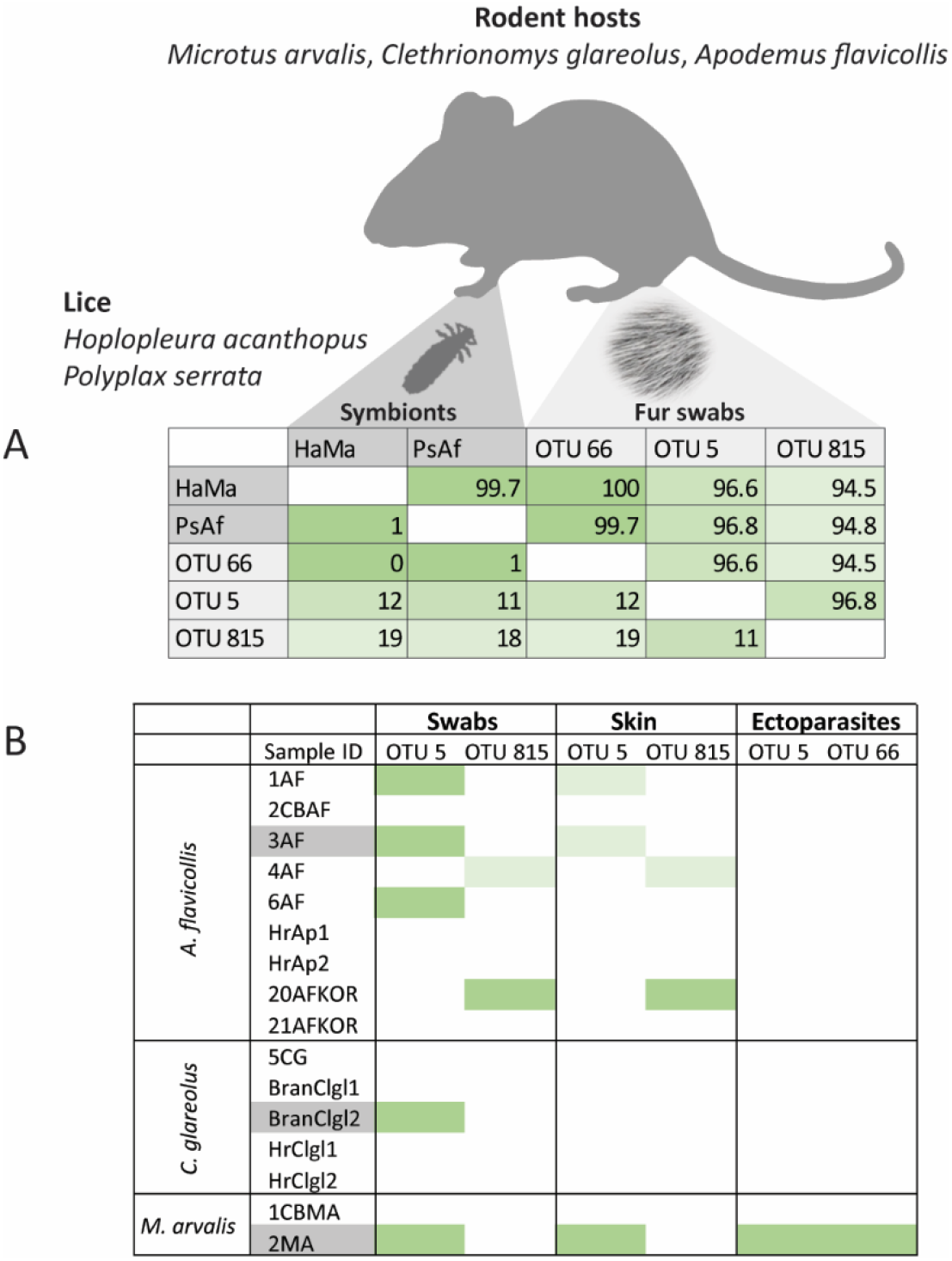
Overview of the Neisseriaceae OTUs determined by amplicon analysis. A - % similarities (above diagonal) and single nucleotide differences (under diagonal) for the 16S rRNA gene fragments acquired from the amplicon screening. HaMa = symbiont of *H. acanthopus*, PsAf = symbiont of *P. serrata*. B - Relative abundance of the Neisseriaceae OTUs comprising more than 10% (light green) or 50% (dark green) of all reads in the sample. Names of the samples chosen for the whole-genome sequencing are highlighted in grey. The samples from mouse spleens are not included in this overview since they did not contain relevant numbers of the reads of the analyzed Neisseraiceae OTUs (see Supplementary Table 1).

### Phylogenetic relationships of the environmental and symbiotic bacteria

In the phylogenomic tree based on a 50-protein matrix, all three specimens of the dominant OTU 5 (for which we obtained a draft genome) and the known Neisseriaceae-related symbionts form a monophyletic clade that shares a relatively long common branch. This arrangement was retrieved by both BI and ML analyses, with only a minor difference in position of *Vitreoscilla stercoraria*, most likely due to extremely short internal branches in this part of the tree (Fig. 2 and Supplementary Figure 1). The shared long branch indicates that the bacteria within the clade diversified only after their common ancestor underwent considerable genome evolution, and its genome became highly different from the other Neisseriaceae. A possible evolutionary scenario based on this phylogenetic arrangement is depicted in Fig. 3. The bacteria from this cluster became part of the rodents’ fur microbiomes, and subsequently established several independent symbiotic relationships with different louse species, some of them evolving into obligate mutualists.

**Figure 2.**
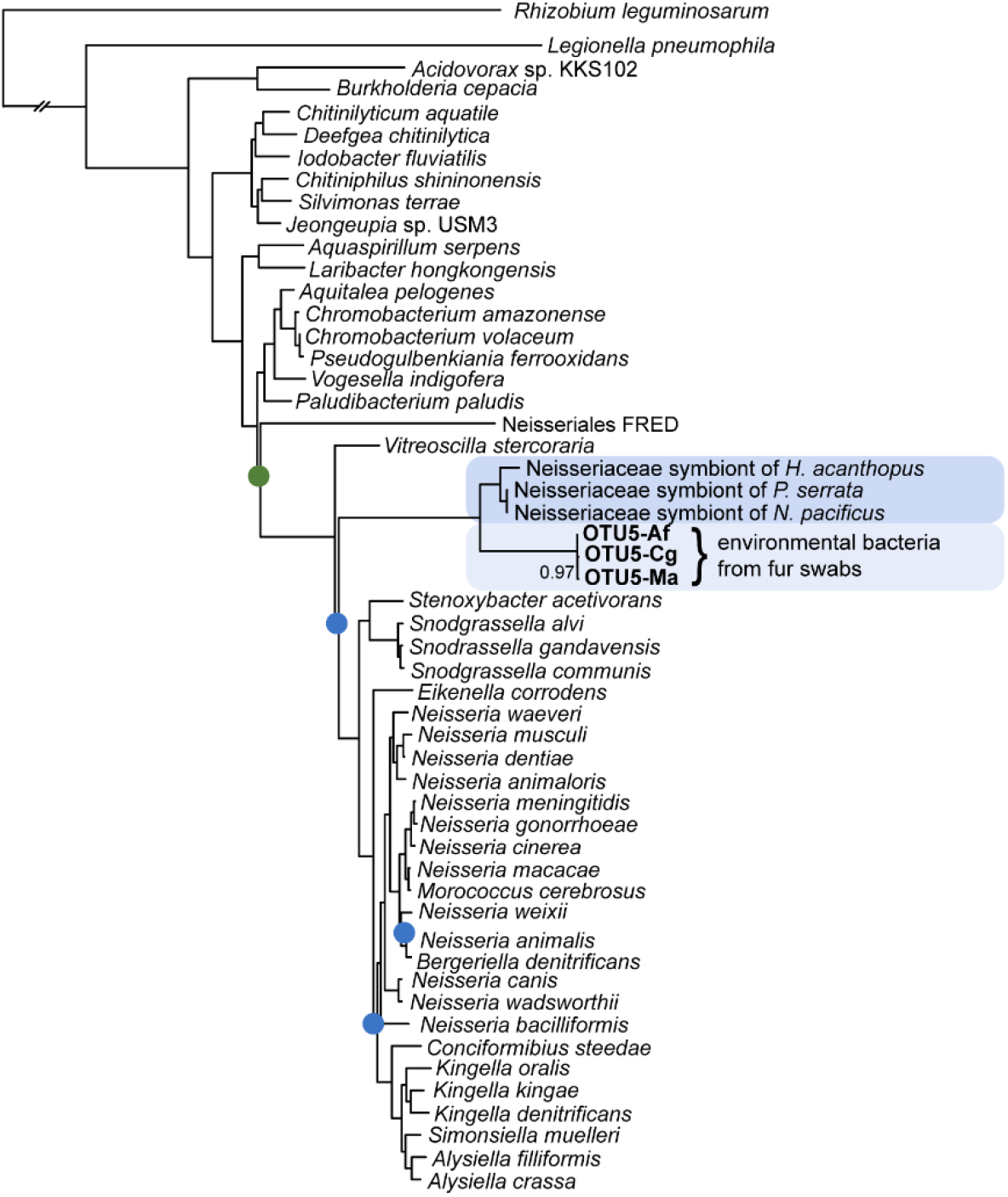
Phylogenetic tree inferred by PhyloBayes from concatenated 50-protein matrix (11,697 aa) using CAT-GTR model. Nodes with posterior probabilities not equal to 1 are designated by the green (0.5) and blue (>0.8) dots. Clustering of the louse symbionts and the environmental OTU 5 samples is highlighted by blue background (dark for symbionts, light for the

**Figure 3.**
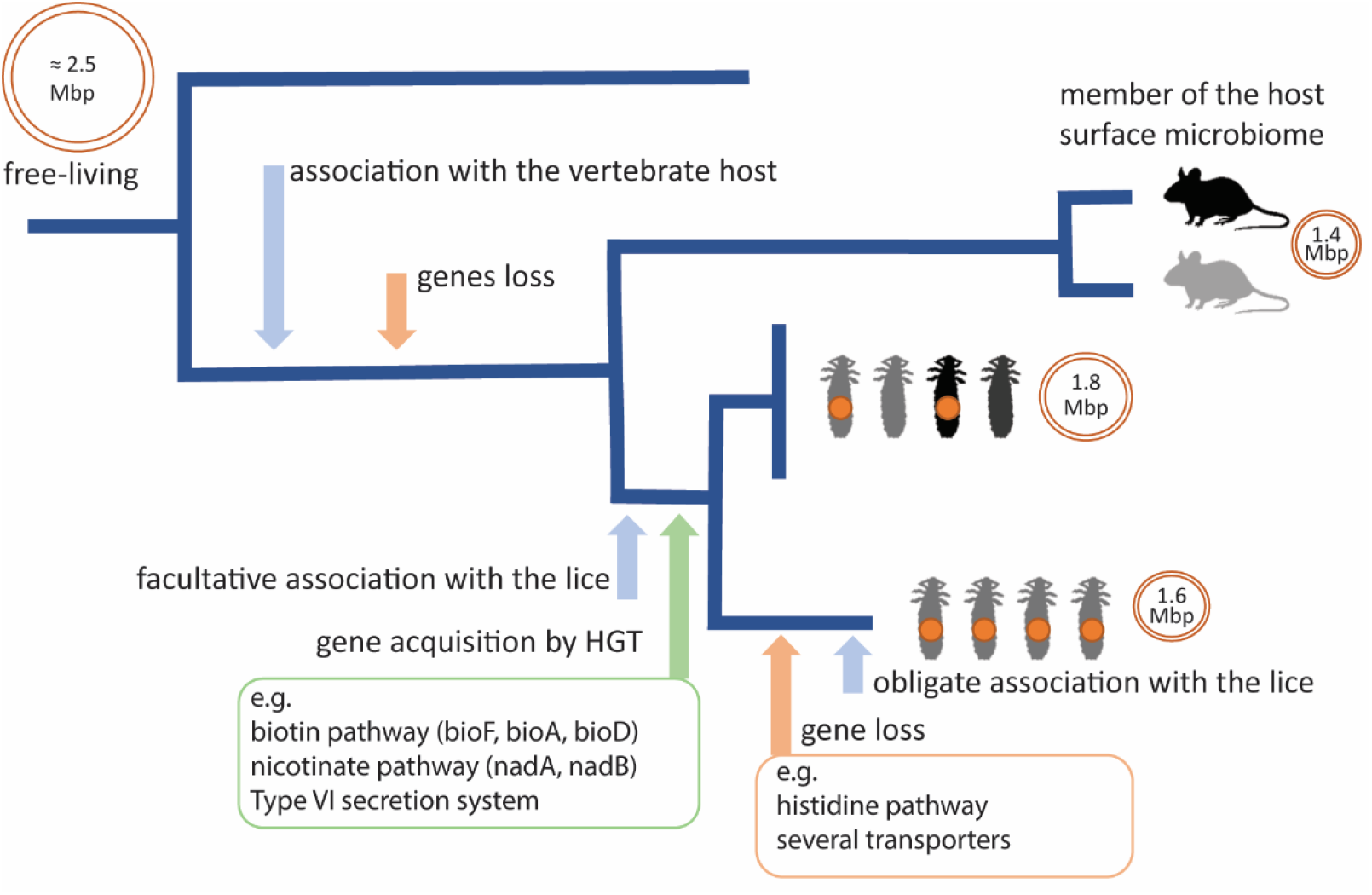
Hypothetical scenario of evolutionary relationship between the microbiome of rodent fur (specifically OTU 5), and the facultative/obligate symbionts in the rodent louse. For details on the gene losses and acquisitions see text.

The phylogeny inferred form the fragment of 16S rRNA gene (which allowed inclusion of a larger taxonomic sample) makes the picture more complex and less straightforward to interpret. It confirms the identity of OTU 66 with the Neisseriaceae related lice symbionts (reported from *H. acanthopus, P. serrata*, and *N. pacificus*), grouping all these symbionts within a “Symbiotic cluster“, together with other ectoparasite-associated bacteria, namely from the boar louse *Haematopinus apri* and the dog prairie flea *Oropsylla hirsuta* (Fig. 4). The OTU 5 branches at the base of a large group (the “OTU 5 cluster”). Apart from several OTUs isolated in this study (OTU 5, OTU 815, OTU 102) it contains an additional 2,876 sequences of bacteria from different environments, obtained from NCBI (Supplementary Table 2).

**Figure 4.**
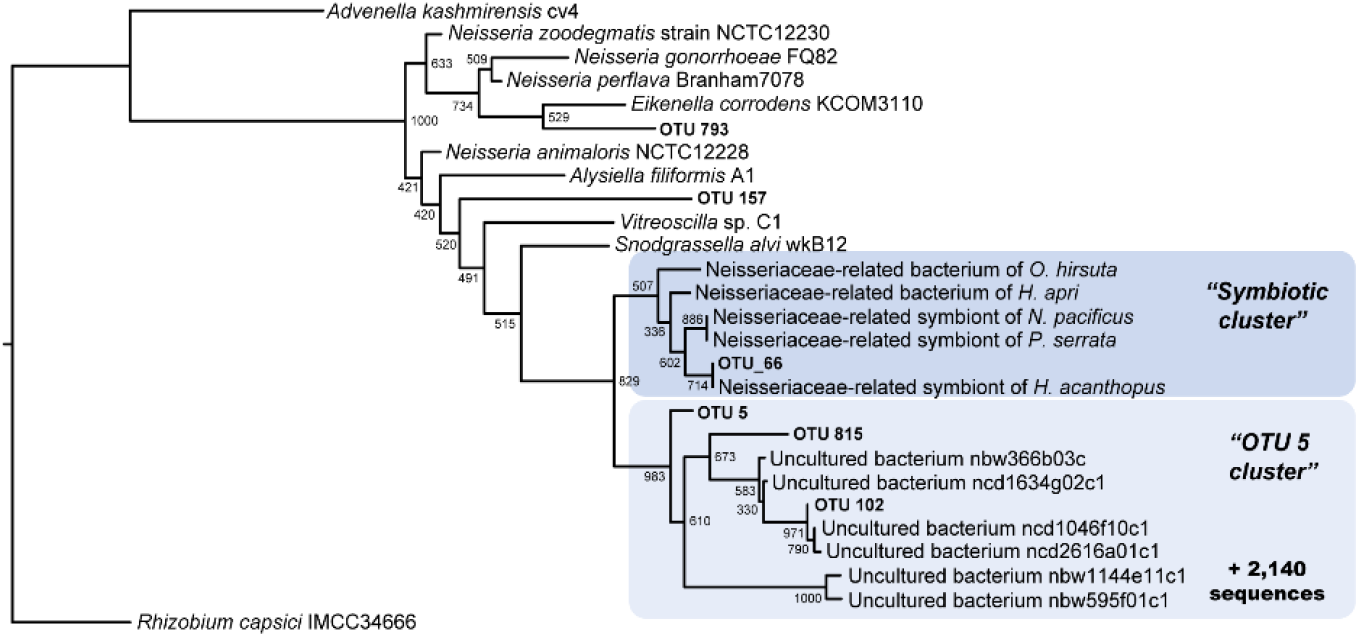
Phylogenetic tree inferred by PhyML analysis from the matrix of 16SrRNA fragments corresponding to the amplicon screening (1,517 bp; ML under HKY85+G+I model). Symbiotic bacteria of lice (“Symbiotic cluster”; designated by dark blue) form a monophyletic lineage together with the Neisseriaceae environmental bacteria (“OTU 5 cluster”; designated by light blue). OTU 66 corresponds to the symbiont of *H. acanthopus*. Values at the nodes show bootstrap supports.

### Genomic comparison

The OTU 5, as the closest relative to the symbiotic cluster, and the dominant constituent of the hosts’ surface microbiome (Fig. 1), is potentially the closest representative of the environmental source which can potentially give rise to the symbionts. We therefore assembled genomes of three samples of OTU 5 obtained from different rodent hosts and compared them to the symbionts described from *Hoplopleura* and *Polyplax* lice (Rihova et al 2021). The metagenomic assemblies of these samples contained fragmented genomes of OTU 5, composed of 9, 34 and 112 contigs (from *M. arvalis, A. flavicollis*, and *C. glareolus*, respectively). The fragmentation of the best draft (OTU5-Ma) was largely due to the rRNA genes. Both 16S and 23S/5S rRNA regions are present in two copies, but due to their identity/high similarity the two copies were assembled as a single contig with twofold coverage compared to the other contigs (contig 6 for 23S/5S rRNA, and contig 9 for 16S rRNA). Compared to the two previously described symbionts of *Hoplopleura* and *Polyplax*, these environmental bacteria (OTU5-Ma, OTU5-Af, OTU5-Cg) possess smaller genomes, but significantly higher GC content. The annotation of these five genomes by PROKKA produced a range of 1225 to 1757 protein-coding genes (Table 1).

When considered in the context of the evolutionary scenario proposed in Fig. 3, the genomic comparison raises an interesting question. The general view predicts that during evolution towards symbiosis, bacteria undergo a typical process of genome degeneration. Its main features are genome shrinking and decrease of GC content (Moran et al 2008). Our data match the latter expectation: the GC content of OTU 5 is consistent with (and provides corroboration of) its free-living/nonsymbiotic nature, compared to the much lower GC content in the two louse symbionts. However, the ratio of sizes seems to contradict the general “paradigm”, as the genomes of the environmental OTU 5 are smaller than those of the symbionts and other free-living Neisseriaceae (which span from 1.8 to 4.3 Mbp). Although we were not able to close the genomes of the environmental OTU 5 into a complete circular chromosome, several lines of evidence indicate that the drafts represent complete or nearly complete genomes. First, even though the three drafts (OTU5-Ma, OTU-Af, OTU5-Cg) were assembled from independent samples, they converge to closely similar sizes and gene contents (Table 1, Supplementary Table 3). Second, although there are additional contigs in both assemblies which were assigned by BLAST also to Neisseriaceae, they differ considerably in their coverage from the OTU 5 contigs. This is particularly well demonstrated in contigs of OTU-Ma where the coverage difference exceeds 1000×. Moreover, these other Neisseriaceae contigs contain genes homologous to those from the OTU 5 contigs and their phylogenetic comparison clearly indicates that they originated from different species (Supplementary Figure 2).

We therefore consider the sizes of the three OTU 5 genome drafts to correspond to the real genome size of this environmental bacterium. While the occurrence of such small genomes is surprising for nonsymbiotic bacteria, it is not entirely unprecedented. For example, among Proteobacteria, several complete genomes of Candidatus *Pelagibacter* sp. (accession numbers GCA_002722165.1, GCA_000195085.1, GCA_000012345.1, GCA_007833635.1, GCA_900177485.1, GCA_012276695.1, GCA_002101295.1) reach sizes between 1.20 to 1.38 Mbp. More interestingly, in respect to the bacterial taxa analyzed here, even smaller complete genome of 0.91 Mbp has been recently reported for a Neisseriales bacterium isolated from an active sludge (Singleton et al 2021; CP064990.1; referred as Neisseriales FRED in the Table 1) . An example, relevant from the ecological point of view, is the bacterium *Dichelobacter nodosus*, living on skin of ungulates, potentially causing dermatitis, which 1.39 Mbp genome (Myers et al 2007; GCA_000015345.1) is close in size to those of OTU 5.

A possible explanation for this seeming inconsistency in the genome sizes can be derived from the blast-based screening which indicated that a considerable proportion of genes in the analyzed genomes are not inherited from the common Neisseriales ancestor but acquired by horizontal gene transfer (HGT). The highest proportion of HGT acquired genes was detected in the facultative symbiont of *P. serrata*, while all other genomes possess smaller and very similar proportions (Fig. 5). The apparently higher level of HGT in the facultative symbiont of *P. serrata* could have led to the genome expansion after the acquisition of the environmental bacterium as a new symbiont (Fig. 3). It is interesting to compare this hypothesis with the observation reported by (Siozios et al 2023) for insect symbionts of the genus *Arsenophonus*. Based on analyses of 38 strains, they concluded that recently acquired symbionts possess larger genomes than their closely related environmentally acquired strains, due to accumulation of horizontally transferred genes.

**Figure 5.**
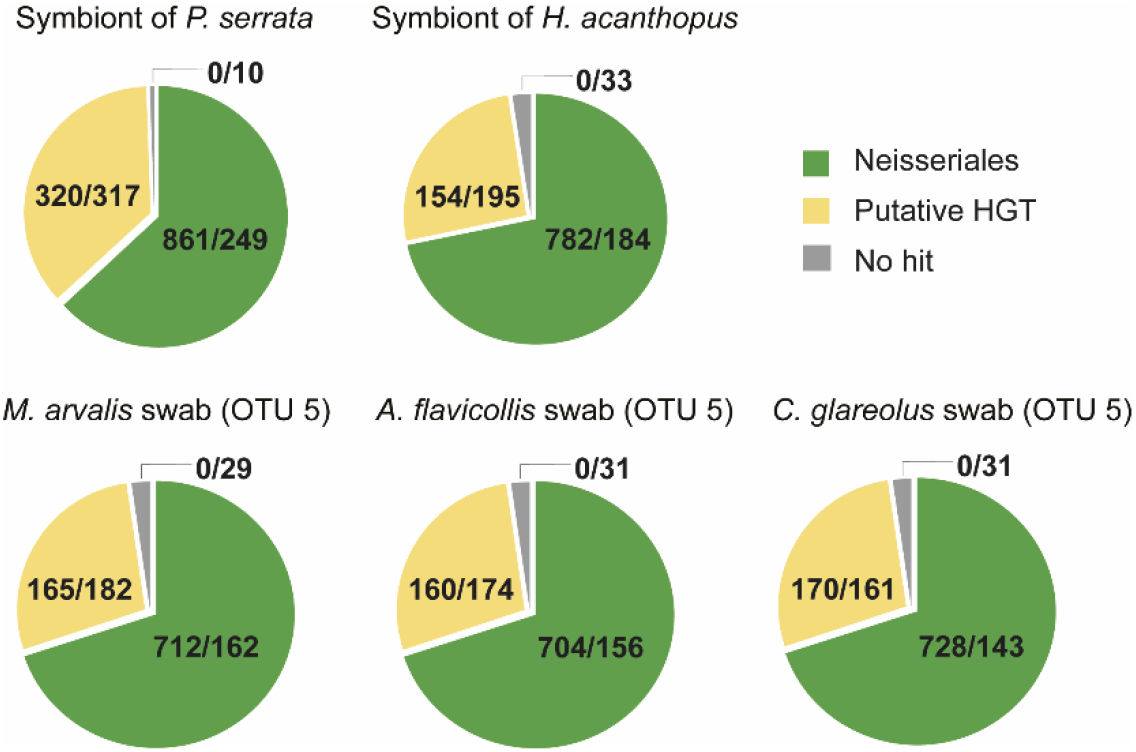
Proportions of the genes of Neisseriales origin (green), the genes acquired by horizontal transfer (yellow), and the genes yielding no blast hits (grey). The numbers stand for the annotated genes/ genes without annotation.

### Metabolic capacities in the symbionts and the environmental bacteria

In correspondence with their genome size, OTU 5 and Neisseriaceae-related symbionts have considerably limited metabolic capacities. Their comparison shows that the five genomes share a large proportion of genes (777) with assigned K numbers (Fig. 6; Supplementary Table 3). Corresponding to its nature of a “young” facultative symbiont with less reduced genome, the *P. serrata* symbiont was the only bacterium with a high number of unique genes (158). Many of them contribute to important metabolic pathways, such as biosynthesis of vitamins (horizontally acquired *nadA, nadB*, and *nadC* in nictotinate pathway), amino acids (*hisA, hisB, hisC, hisD, hisF, hisI, hisG, hisH*, and *hisZ* in histidine), metabolite transporters, or components of secretion systems. In the obligate symbiont HaMa with stronger genome reduction, only eleven genes were identified as unique, encompassing e.g., parts of secretion system or DNA replication and repair (Supplementary Table 4).

**Figure 6.**
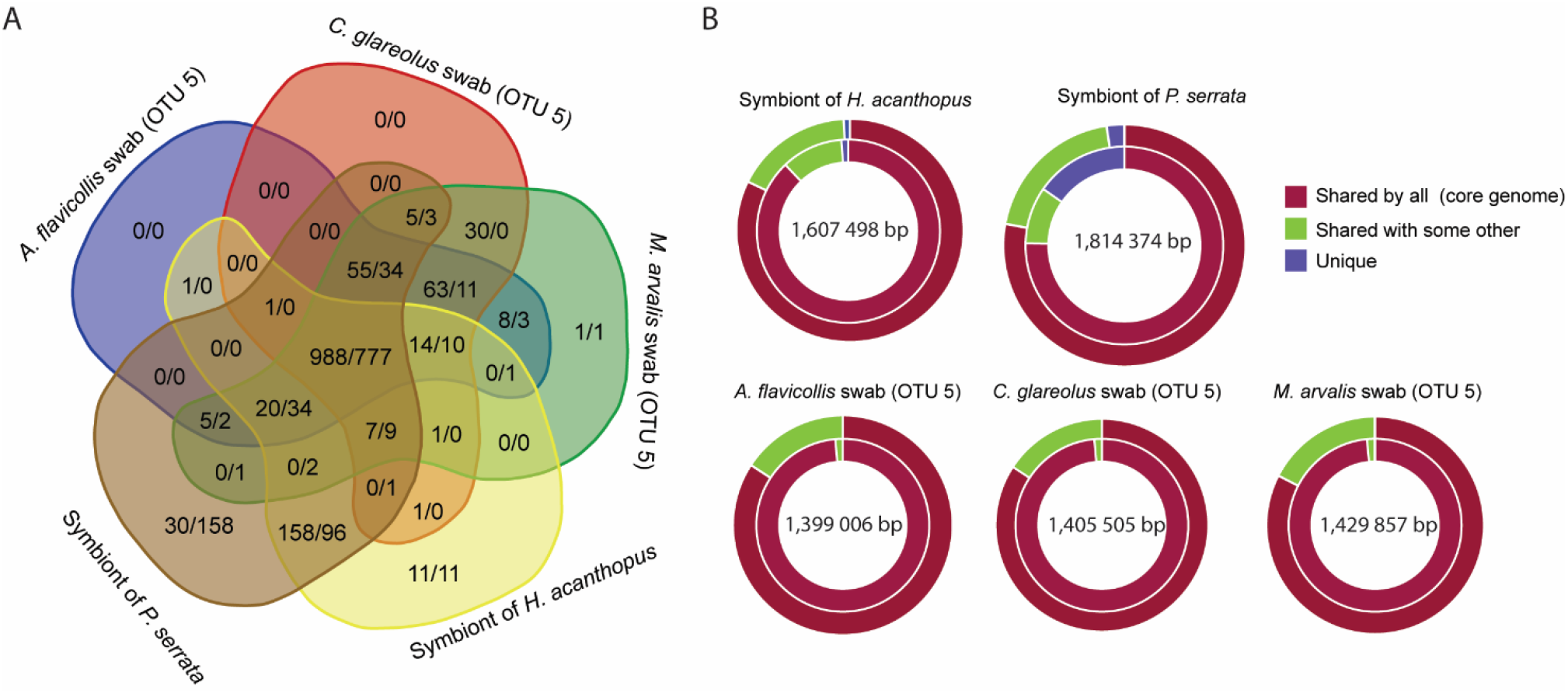
Comparison of genome contents of the Neisseriaceae related symbionts and the environmental bacteria. A – Number of genes shared among the genomes. The numbers stand for orthologs/genes with assigned K numbers. B – Proportion of the shared genes: outer circles show orthologs, inner circles the genes with assigned K numbers.

To assess the genomes in context of the hypothetical evolutionary scenario (Fig. 3), we performed a detailed comparison which also included three additional Neisseriaceae. Two species are closely related to the “symbiotic” and “OTU 5” clusters (*Snodgrassella alvi* and *Vitreoscilla stercoraria*), and the third, *Neisseria cinerea*, is a Neisseriaceae free-living species with small genome (1.7 Mbp). Reconstruction of selected metabolic capacities and other cellular functions for these seven genomes is summarized in Supplementary Table 4. Two metabolite categories are often discussed in connection to bacterial symbionts in insects, amino acids, and B vitamins and cofactors. For amino acids, our comparison does not indicate any differences related to different lifestyles of the OTU 5 and the two symbionts. Both types are able to synthesize a considerable number of amino acids, including proline, lysine, cysteine, valine, isoleucine, leucine, tryptophan, threonine, arginine, and glutamine (Supplementary Table 4). A different picture is, however, provided by comparison of B vitamins, where two pathways, biotin, and nicotinate, differ between the environmental OTU 5 and the symbionts.

In the biotin pathway, all compared genomes except *N. cinerea* seem to lack the “original” (i.e. gene homologs directly inherited from the Neisseriaceae ancestor) of the central genes bioF, bioA, bioD, and bioB. In the two symbiotic bacteria, these essential genes are present, but at least three of them have been acquired by HGT (bioF, bioA, and bioD, while bioB produces an inconclusive BLAST result; Fig. 7, Supplementary Table 5). A possible explanation is that the biotin synthesis capacity was lost by the common ancestor of OTU 5 and the symbionts, and acquired secondarily by the ancestor of the symbionts, perhaps in relation to its symbiotic function. The only missing gene in the symbionts (bioH) has been shown in several studies as replaceable by various other genes (Bi et al 2016, Hang et al 2019). The presence of nine (out of the ten) required genes indicates that the biotin pathway is indeed functional in the two symbionts, and the vitamin is likely provided to the host. For the nicotinate pathway the situation is similar, but more complex. All four genomes (OTU 5 and the symbionts) lack Neisseriaceae homologues of three central genes (nadB, nadA, and nadC). Their horizontally acquired homologues are only present in the facultative symbiont of *P. serrata*. It is however difficult to speculate whether this HGT occurred only in the *P. serrata* symbiont, or in the common ancestor of both *P. serrata* and *H. acanthopus* symbionts, and the genes were then lost during the genome economization of the latter.

**Figure 7.**
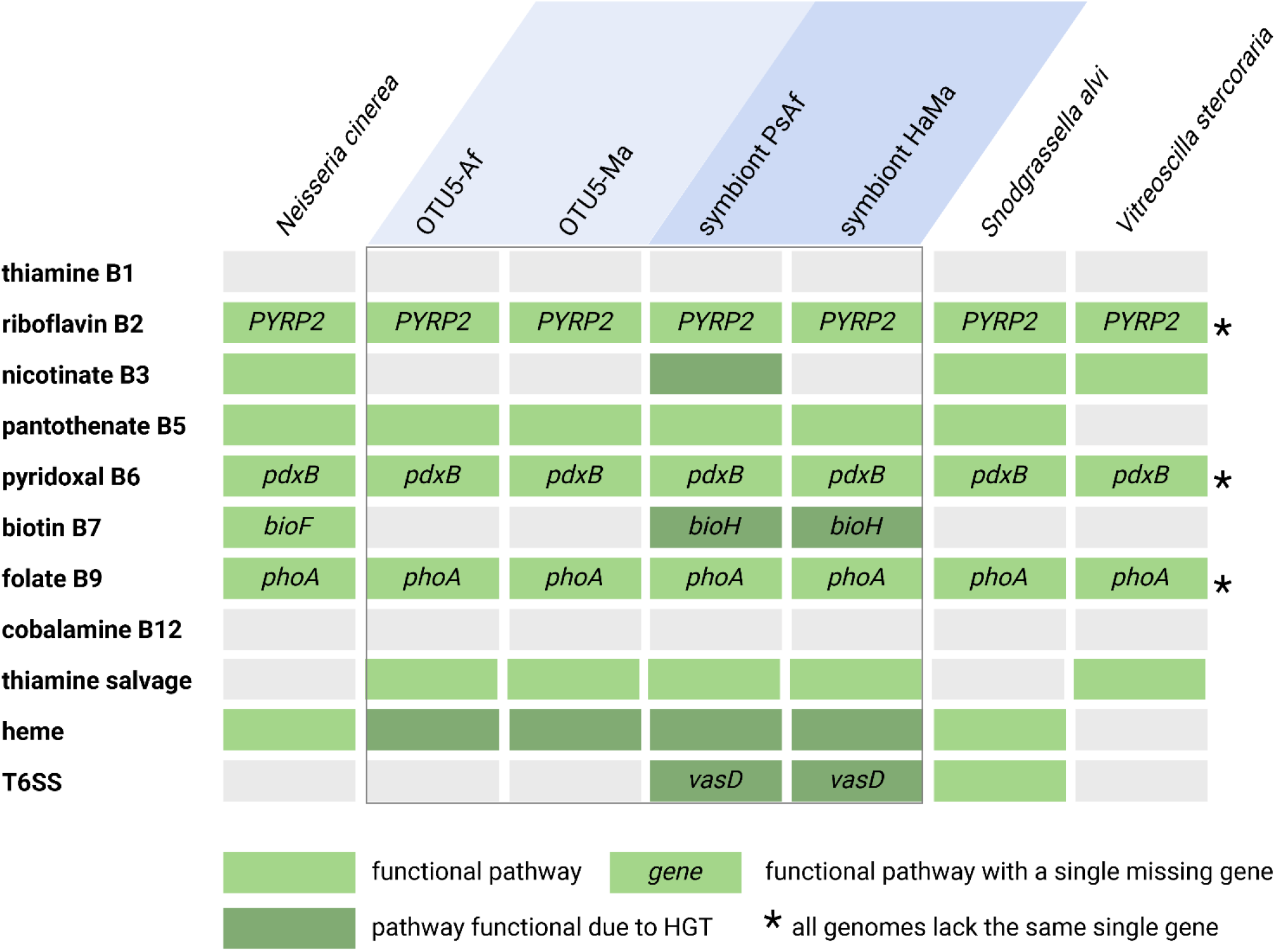
Comparison of selected metabolic capacities of environmental OTU 5 (light blue background), louse symbionts (dark blue background), and three related Neiseriaceae. Colours of the cells stand for: grey - missing/nonfunctional pathways, light green - functional pathway, dark green – pathway functional due to HGT. Names in the cells indicate missing genes, which are however likely not essential for the pathway function (see the text). Asterisks indicate pathway with the same single gene missing in all genomes. Following genes are putative HGT in the dark green cells: nicotinate - nadB, nadA, nadC; biotin - bioF, bioA, bioD, bioB; heme - ALA5; T6SS - vasK, vasF, vasE, vasB, vasA, impF, impC, impB, impA, hcp, vgrG, vasG. For complete metabolic overview see Supplementary Table 4.

The remaining B vitamin pathways do not show differences between OTU 5 and the symbionts. Two of them miss several genes and are obviously nonfunctional in all compared genomes (thiamine and cobalamin). They however possess the three genes composing thiamine salvage pathway. The loss of thiamine synthesis has been previously observed in other symbionts of bloodsucking insects (Boyd et al 2017, Martin Rihova et al 2023, Rihova et al 2017), indicating that this vitamin may be obtained from the vertebrate host blood.

Finally, the remaining four pathways seem to be functional in both the OTU 5 genomes and the symbionts (pantothenate, riboflavin, pyridoxal, and folate). However, while for pantothenate these bacteria possess all required genes, for the latter three pathways the situation differs. All seven compared genomes lack the same single gene in an otherwise complete pathway (based on KEGG definitions as reference). A similar situation was also detected in several other pathways presented in the Supplementary Table 4. As stated by Rihova et al 2021, such cases indicate that the pathways are likely functional, and the “missing” genes can be either not essential or their role can be substituted with different genes.

Similar to biotin discussed above, the pattern of the type VI secretion system (T6SS) seems to also match the presumed evolution of the OTU 5 + the symbionts cluster. Generally, secretion systems are considered important devices of symbiotic bacteria in particular stages of their evolution. In our analysis, both OTU 5 and the two louse symbionts lacked completely T1SS, T2SS, and T3SS. Of the T4SS, they possessed various numbers of the vir genes. The most interesting, however, was the pattern for the T6SS, a system known from various bacteria, including symbionts (Guckes and Miyashiro 2023, Suria et al 2022, Takeshita and Kikuchi 2020), which is often subject to HGT. This system was absent in the environmental OTU 5 bacteria, but present in both louse symbionts where it was obtained secondarily by HGT. The only missing gene, vasD (tssJ), was previously shown as likely not essential for the T6SS function (Suria et al 2022). Interestingly, the only other bacterium with T6SS was *S. alvi*, living also as an insect symbiont.

In summary, the close phylogenetic relationships, and genomic similarity, between the symbionts of several louse species and the environmental bacteria identified in fur of the rodent hosts, suggest that the fur microbiome can serve as a source of louse symbionts. Since these Neisseriaceae-related symbionts have been found in several phylogenetically distant louse species, it is obvious that they established their symbiosis in several independent events. This further supports the view on the fur microbiome as a background for the evolutionary origin of lice-bacteria symbiosis.

## Methods

### Sampling design and DNA extraction

The sample was collected in April 2021 from 16 mice (*Apodemus flavicollis* 9, *Clethrionomys glareolus* 5, *Microtus arvalis* 2) from either death snap traps or Sherman live traps. The animals collected from live traps were euthanized before dissection. From each mouse we sterilely sampled spleen, skin (piece from mouse hip), surface swab from mouse posterior ridge, and ectoparasites, and stored these in UV-sterilized absolute ethanol. DNA isolates from swabs and ectoparasites were extracted using the QIAamp DNA Micro Kit (Qiagen). For DNA extraction from skin pieces and spleens we used DNeasy Blood & Tissue Kit (Qiagen).

### Amplicon library preparation and sequencing

DNA templates were processed for the 16S rRNA gene amplification according to Earth Microbiome Project standards (EMP; http://www.earthmicrobiome.org/protocols-and-standards/16s/). Samples were multiplexed based on a double barcoding strategy with 12-bp Golay barcodes (EMP protocol) combined with the forward primer 515F (Parada et al 2016) and 926R primers (Parada et al 2016, Walters et al 2016) yielding 450 bp long amplicons of V4 hypervariable region of the 16S rRNA gene. The library was sequenced on the single MiSeq run using v2 chemistry. To confirm the barcoding output and evaluate any possible amplification bias and the sequencing depth, we used one sample of mock communities with an equal composition of 10 bacterial species and one sample with a staggered composition of the same 10 bacteria (ATCC Microbiome Standards). The output was set to 2 × 250 bp (paired-end reads). We used 2 negative controls to assess the extraction and amplification procedures, i.e., control for extraction procedure and PCR water templates. The metadata for host species, sampling localities, accession numbers and number of ectoparasites are presented in Supplementary Table 1. Raw amplicon reads were deposited in the NCBI Sequence Read Archive (SRA) repository under BioProject accession number PRJNA1086898.

### Amplicon library processing and analyses of the amplicons

The raw reads were quality checked using FastQC (Andrews 2010) and trimmed using USEARCH v9.2.64 (Edgar 2013). The reads were further processed into operational taxonomic units (OTUs) with in-house workflow implementing suitable scripts (Brown et al 2020, Rodriguez-Ruano et al 2020) from USEARCH. The dataset was then clustered at 100% identity providing a representative set of sequences for *de novo* OTU picking. OTUs were defined at 97% identity match using the USEARCH global alignment option. Taxonomical assignment to the representative sequences was assessed using the best BLAST hits (Camacho et al 2009) against SILVA 132 database (Quast et al 2013). Mitochondrial OTUs, chloroplast sequences, singletons and extremely low abundant OTUs were removed from the final OTU table using QIIME 1.9 (Caporaso et al 2010). The final OTU table was filtered for potential contaminants presented in the negative controls (Supplementary Table 1).

### 16S rRNA gene phylogeny

To place the dominant OTU 5 bacterium within a broad sample of Neisseriaceae, we screened 16S rRNA genes available in the NCBI GenBank (https://www.ncbi.nlm.nih.gov/). We used the sequence of OTU 5 as a query for discontinuous MegaBLAST (Morgulis et al 2008) set to 3,000 hits (E-value 0.05). Of the resulting hits, 80 sequences corresponded to 19 complete genomes (due to multiple copies of the 16S rRNA genes) and the remaining 2,920 sequences were complete or partial 16S rRNA genes. To build phylogenetic matrix, we used a single arbitrarily chosen copy to represent each of the 19 complete genomes. The resulting 2,937 sequences were filtered for duplicates, and only unique sequences were retained (2,884 sequences). The vast majority of these sequences originated from human samples (2,802), mostly from skin (2,718). The rest of the sequences represented bacteria with a large variety of origins, such as air, sediment, and dust samples (Supplementary Table 2). To set the analysis in a broader phylogenetic span, and to provide a reliable outgroup, we further included 16S rRNA genes of several additional bacteria (Supplementary Table 2). Together with the six Neisseriaceae OTUs (5, 66, 102, 157, 793, and 815), the Neisseriaceae-related bacteria described from lice and flea (Jones et al 2008, Nishide et al 2022), and ten outgroups, the final matrix contained 2,905 sequences (see Supplementary Table 2 for detailed description of the matrix composition). These sequences were aligned using MAFFT with E-INS-i settings (Katoh et al 2002) implemented in Geneious Prime v.2020.2.5 (Kearse et al 2012). The alignment was processed in Gblocks using less stringent options. The length of the resulting matrix was 1,035 bp. The phylogenetic reconstruction was performed using the ML method in IQ-TREE 2 (Minh et al 2020) with an automatic selection of the model (GTR+F+R5 chosen according to BIC).

Apart from this large 16S rRNA gene tree, we inferred a reduced 27-taxa tree, which allowed for a clear visualization of the Neisseriaceae OTUs positions within the whole Neisseriaceae family (see list of used sequences in Supplementary Table 2). The taxa selected for this tree were realigned by MAFFT v.7.450 (with E-INS-i settings) in Geneious Prime. The size of this reduced matrix was 1,517 bp. The phylogenetic tree was inferred using the ML method using online PhyML 3.0 under HKY85+G+I model chosen using SMS function of online PhyML with 1000 bootstrap replicates. All matrices and phylogenetic tress are deposited in Mendeley Data under the “doi” link 10.17632/3wv3szky46.2.

### Genome assembly

Based on the amplicon analysis and the phylogeny derived from 16S rRNA gene, we selected for the metagenomic assembly three samples which contained Neisseriaceae OTU 5 as the most abundant bacterium (3AF from *Apodemus flavicollis*, 2MA from *Microtus arvalis* and BranClgl2 from *Clethrionomys glareolus*; see Fig. 1). We measured DNA concentration using the Qubit High sensitivity kit and assessed the DNA quality by gel electrophoresis. DNA was sequenced on one Sp lane of Illumina NovaSeq6000 platform (W. M. Keck Center, University of Illinois at Urbana-Champaign, Illinois, USA) using 2 × 250 paired-end reads. The reads were checked by FastQC and trimmed using the BBtools package with settings minlen=240 and maq=20 (https://jgi.doe.gov/data-andtools/bbtools). The resulting dataset contained 116,729,096 reads (2MA), 94,146,136 reads (3AF) and 182,717,424 reads (BranClgl2). Assemblies were done in SPAdes v.3.15.2 (Bankevich et al 2012) with --meta option. To identify the contigs which potentially represent the Neisseriaceae genome(s), we employed the following complex procedure. We first screened the complete assemblies by BLASTn (Altschul et al 1990) using genes from two Neisseriaceae-related louse symbionts (CP046107 and WNLJ00000000), *Vitreoscilla* sp. C1 (CP019644.2), *Snodgrassella alvi* wkB2 (GCF_000600005.1), *Neisseria gonnorrhoeae* (GCA_013030075.1), *Neisseria animaloris* (GCF_002108605.1), *Eikenella corrodens* (GCF_000158615.1), and *Alysiella filiformis* (GCA_014054525.1) as queries (E-value set to 10.0 and hit number to 1). This approach ensured that each candidate contig was identified multiple times. The resulting sets of the identified contigs (34 contigs for the sample 2MA, 59 contigs for 3AF, and 168 contigs for BranClgl2) contained both prokaryotic and eukaryotic contigs with genes yielding significant BLAST hits. Eukaryotic contigs were removed manually based on the ORFs density as predicted in Geneious Prime. A more precise taxonomical assignment of the remaining contigs was done by BLASTn search (E-value 1, three best hits) against the GenBank nr database. The contigs with at least one Neisseriaceae hit were considered as potentially representing Neisseriaceae OTUs and used in the following steps.

Since the microbiomes of each sample contained several different Neisseriaceae OTUs, we further filtered the contigs with the aim to extract those corresponding to OTU 5. We used two criteria to verify the identification of the contigs. First, since different OTUs occurred in the samples with different abundances (based on amplicon analysis), their contigs possessed different ranges of coverage. To obtain contigs corresponding to the most abundant OTU 5, we extracted sets of contigs with the highest coverage (app. 1000x for 2MA, 8x for 3AF and 8x for BranClgl2). Second, as an additional confirmation, we compared gene content of the high-coverage contigs with the remaining low-coverage contigs (see below for annotation procedure). This comparison revealed frequent gene duplicates (or even triplicates), with one copy present in the high-coverage contig and the other copy in the low-coverage contig. To confirm that the contigs carrying these gene duplicates, but assembled with different coverages, belong to phylogenetically different bacteria, we arbitrarily chose one gene from each of these contigs and reconstructed their phylogenetic relationships in Neissericeae context (Supplementary Figure 2). The trees were inferred by maximum-likelihood in PhyML (Guindon et al 2010) as implemented in Geneious Prime. The sets of the high-coverage contigs were considered as draft genomes of the OTU 5, composed of 9 contigs from 2MA (creating the genome draft OTU5-Ma), 34 contigs for 3AF (draft OTU5-Af), and 112 contigs for BranClgl2 (draft OTU5-Cg). The SRA data and corresponding draft genomes were deposited in GenBank under the BioProject accession number PRJNA1086898. Annotations for these draft genomes are deposited in Mendeley Data under the “doi” link 10.17632/3wv3szky46.2. All three genome drafts were annotated using PROKKA (Seemann 2014) and used for the downstream analyses. Based on the annotations, we inferred the main characteristics of the new genome drafts and their relatives (coding density, number of genes, mobile elements, etc.; Table 1). The number of phages was further assessed by PHASTER (Arndt et al 2016), and potential pseudogenes were identified by Pseudofinder (Syberg-Olsen et al 2022). The completeness of the genome drafts was evaluated using BUSCO v4.0.6 (Simao et al 2015). The overall similarity of the OTU 5 samples was measured by the average nucleotide identity (ANI) using the online ANI calculator (Yoon et al 2017).

### Genome comparison

We compared three different genomic traits of the environmental OTU5 bacteria and the Neisseriaceae-related symbionts. First, we compared numbers of the PROKKA-annotated genes, shared orthologs identified by OrthoFinder (Emms and Kelly 2019), and shared genes with K numbers assigned by BlastKOALA server (Kanehisha et al 2016). Second, to compare the proportions of genes obtained by HGT, we performed taxonomical assignment of each annotated gene using BLASTp search against nr database (set to ten hits and E-value 0.1). We applied a conservative approach, with the aim not to overestimate the effect of HGT. The genes which contained at least one member of Neisseriales among the hits were assigned to the category “Neisseriales”. The genes with all hits representing different bacterial groups (no Neisseriales hits) were assigned to the category “Putative HGT”. The rest of the genes returned no hits (assigned to the “No hit” category). Third, to assess metabolic capacities of each genome, we mapped the BlastKOALA-assigned functions on metabolic pathways using Kyoto Encyclopedia of Genes and Genomes (KEGG) database. The metabolic capacities of the OTU5 bacteria were compared to several additional bacteria, including the Neisseriaceae-related symbionts, their closest relatives (*Snodgrassella alvi* and *Vitreoscilla stercoraria*), the smallest environmental Neisseriales bacterium for NCBI (Neisseriales FRED) and one common commensal Neisseraceae species (*Neisseria cinerea*). Accession numbers for all compared genomes are listed in Table 1. The K numbers and annotations for compared genomes are deposited in Mendeley Data under the doi link 10.17632/3wv3szky46.2.

### Phylogenomic analyses

To verify phylogenetic positions of the OTU 5 samples in a robust phylogenomic framework, we used the dataset of Neisseriales proteomes included in Rihova et al 2021 and extended it with additional proteomes available from the NCBI database, including the putatively closest relatives (Neisseriaceae-related symbionts of lice). Representatives of other bacterial groups, alphaproteobacterium *Rhizobium leguminosarum*, gammaproteobacterium *Legionella pneumophila*, and two betaproteobacteria from the order Burkholderiales, *Acidovorax* sp. KKS102 and *Burkholderia cepacia*, were used as outgroups (Supplementary Table 2). To build the matrix we selected a set of fifty single-copy orthologs determined by OrthoFinder, for which we were able to reliably establish their origins within Neisseriales (i.e., the orthologues from the BLAST category “Neisseriales”; details on the assignment to the categories are provided above in the section Genome comparison), and which were present in all genomes. The sequences were aligned in MAFFT under E-INS-i settings implemented in Geneious Prime platform and processed by Gblocks 0.91b (Talavera and Castresana 2007) using more stringent options. The resulting matrix consisted of 11,697 amino acids. For the phylogenetic reconstructions we used the maximum-likelihood (ML) and Bayesian inference (BI). The ML tree with 100 bootstrap replicates was generated using IQ-TREE 2 with the best fitting mixture model Q.pfam+C60+F+R4 selected by the ModelFinder implemented in IQ-TREE 2 program. BI was done by Phylobayes (Lartillot et al 2013) under CAT-GTR model. Two independent chains were run for 50,000 generations when their maxdiff reached 0.07 and effective sizes for all parameters exceeded 300 (with burn-in set to 10,000). Used matrices and phylogenetic tress are deposited in Mendeley Data under the “doi” link 10.17632/3wv3szky46.2.

