## Supplementary Figure 1 for "Fur microbiome as a putative source of symbiotic bacteria in sucking lice"

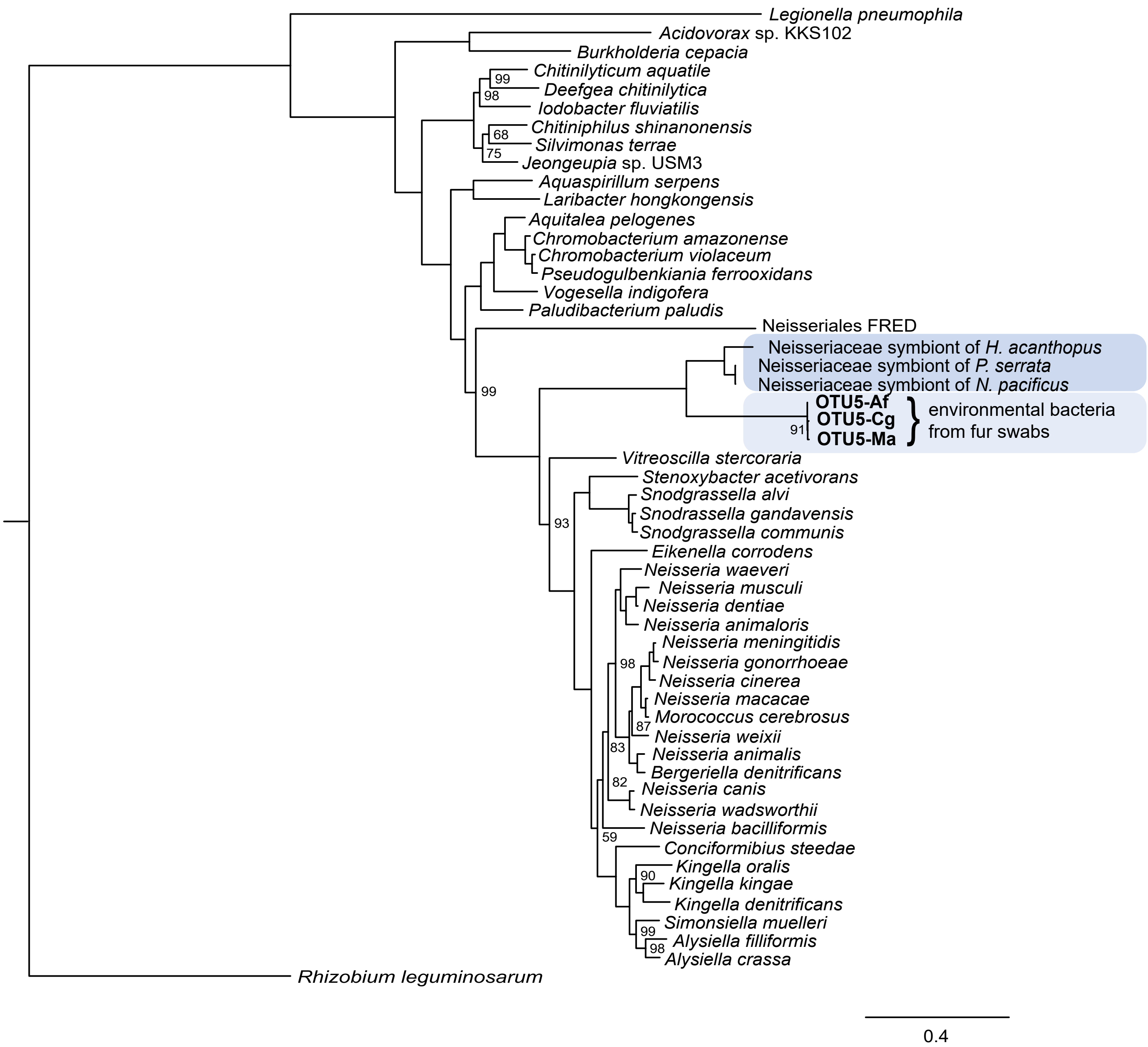


**Supplementary Figure 1.** Phylogenetic tree inferred by IQ-TREE 2 from concatenated 50-protein matrix (11,697 aa) using the best fitting mixture model Q.pfam+C60+F+R4 selected by the ModelFinder implemented in IQ-TREE 2 program. Clustering of the louse symbionts and the environmental OTU 5 samples is highlighted by blue background (dark for symbionts, light for the environmental OTU 5).
