## Supplementary Figure 1 for "Fur microbiome as a putative source of symbiotic bacteria in sucking lice"

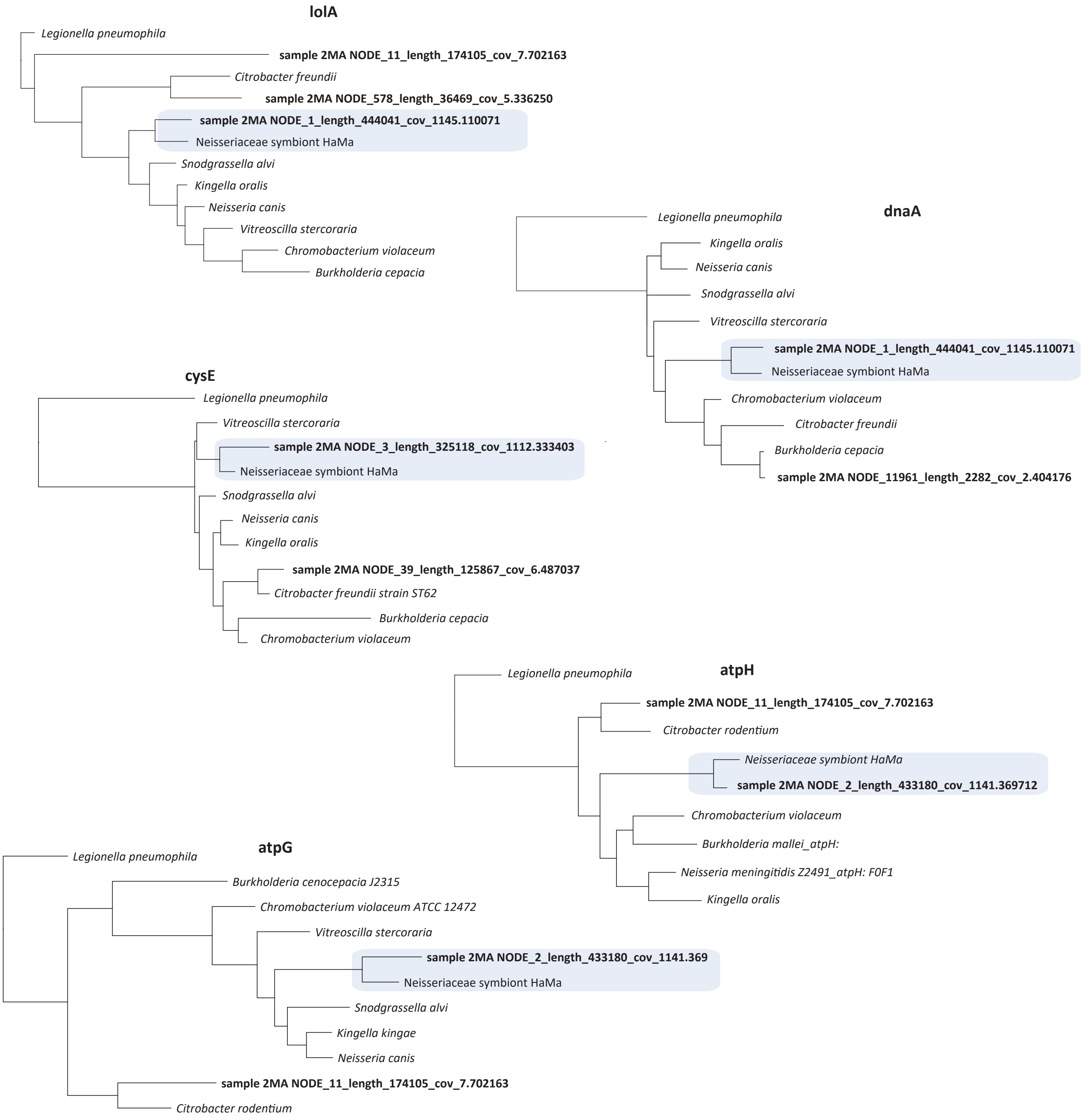


| | lolA | dnaA | cysE | atpH | aptG |
|---|---|---|---|---|---|
| *Burkholderia cenocepacia* | | | | | CP102475 |
| *Burkholderia cepacia* | CP095498 | CP095498 | CP090608 | | |
| *Burkholderia mallei* | | | | CP090608 | |
| *Chromobacterium violaceum* | LR134182 | CP069587 | AE016825 | CP024029 | CP050992 |
| *Citrobacter freundii* | CP140972 | CP140972 | CP099037 | | |
| *Citrobacter rodentium* | | | | CP038008 | CP082833 |
| *Kingella kingae* | | | | | CP050135 |
| *Kingella oralis* | CP059569 | CP059569 | CP059569 | CP059569 | |
| *Legionella pneumophila* | CP021268 | CP114576 | CP115862 | CP045974 | CP115862 |
| *Neisseria canis* | LR134313 | LR134313 | LR134313 | | LR134313 |
| *Neisseria meningitidis* | | | | CP045961 | |
| *Snodgrassella alvi* | CP132376 | CP132374 | CP132375 | | CP132375 |
| *Vitreoscilla stercoraria* | CP091512 | CP091513 | CP091514 | | CP091516 |

**Supplementary Figure 2.** Phylogenetic analyses of duplicated genes in the assembly of the 2MA sample. The genes from the 2MA assembly are printed in bold, and the description provides contig number, length, and coverage. Accession numbers of the genes used in the matrices are listed in the table. The genes corresponding to the OTU 5 and the closely related symbiont of *Hoplopleura acanthopus* collected from *Microtus arvalis* (HaMa) are designated by blue background in the trees.
